# Exploring the interplay of complex carbohydrate intake, the microbiome CAZymes pool and short-chain fatty acid production in the human gut: insights from different cohorts in the Argentine population

**DOI:** 10.1101/2025.05.15.654276

**Authors:** Milagros Trotta, Camila Agustini, Cristian Rohr, Ricardo Martin Ame, Laureano Giordano, Domingo C. Balderramo, Pablo A. Romagnoli, Martin Vazquez

**Affiliations:** Heritas (www.heritas.com.ar), Argentina; Hospital Privado Universitario de Córdoba, Córdoba, Argentina; Centro de investigación en Medicina Traslacional “Severo R. Amuchastegui” (CIMETSA-CONICET), Instituto Universitario de Ciencias Biomédicas de Córdoba (IUCBC), Córdoba, Argentina; Consejo Nacional de Investigaciones Científicas y Técnicas, CONICET, Argentina

**Keywords:** Gut microbiome, CAZymes, SCFAs, IBD, microbial diversity, enzymatic functionality, microbiome resilience

## Abstract

Carbohydrate-active enzyme (CAZyme) activity in the gut microbiome has significant implications for health, including the release of nutrients otherwise inaccessible to the host and the enhancement of digestive efficiency. The primary end products of indigestible carbohydrate fermentation are short-chain fatty acids (SCFAs). We hypothesized that increased dietary fiber consumption could lead to greater SCFA production in the human gut. To investigate this, we examined the relationship between complex carbohydrate intake, CAZyme activity in the human gut microbiome, and SCFA production at the whole metagenomic level across three cohorts: a healthy reference-controlled cohort, an average cohort of individuals living in industrialized cities, and a cohort of patients with inflammatory bowel disease (IBD). Metagenomic sequencing and bioinformatic analyses were utilized to assess the diversity, abundance, and functionality of CAZymes, as well as the metabolic capacity for SCFA production.

The average cohort exhibited higher alpha diversity of CAZyme families compared to the reference-controlled cohort, although subfamily composition was similar between both. A moderate negative correlation was identified between CAZyme abundance and SCFA production, indicating that a higher number of these enzymes does not directly translate to increased SCFA synthesis. In IBD patients, a decrease in the diversity and composition of CAZyme subfamilies was observed, suggesting a disruption in enzymatic functions associated with the disease. However, the overall functionality of CAZymes remained relatively stable across different health conditions, highlighting the resilience of the gut microbiome for these functions.

These findings deepen our understanding of the gut microbiome’s role in health and disease, emphasizing that despite variations in microbial diversity, key enzymatic functions persist. The study underscores the complexity of the non-linear relationship between complex carbohydrate metabolism and SCFA production, laying the groundwork for future research on microbiome-targeted therapeutic/dietary profile interventions in both non-disease and chronic diseases conditions.

## Introduction

Humans function as superorganisms living in symbiosis with billions of bacteria and eukaryotic cells. This symbiosis between the human host and its microbes is referred to as a “holobiont” (Rosenberg & Zilber-Rosenberg, 2016), while the collective genomes of these organisms are known as the “hologenome”. The holobiont’s ability to adapt and evolve is driven by modifications in both the host’s genome and its microbiome. Among the various microbial communities in the human body, the gut microbiota plays a particularly important role.

The gut microbiota refers to the complex community of microorganisms inhabiting our gastrointestinal tract, which exerts significant influence on the host during both health and disease (Thursby & Juge, 2017). The composition of the gut microbiota includes bacteria (e.g., 99% of the gut microbiota consists of Firmicutes, Bacteroidetes, Proteobacteria, and Actinobacteria), fungi (e.g., Candida), viruses (e.g., bacteriophages), and parasites (e.g., flagellates) (Beam et al., 2021).

One of the primary roles of microorganisms within the gut microbiota is the digestion of complex carbohydrates through specialized enzymes known as Carbohydrate-Active enzymes (CAZymes). These CAZymes, encoded by the gut microbiota, catalyze the breakdown of glycoconjugates, oligosaccharides, and polysaccharides into fermentable monosaccharides. The human genome encodes only 17 glycoside hydrolase enzymes for digesting dietary glycans, specifically starch, sucrose, and lactose (Kaoutari et al., 2013). This limited capacity contrasts with the vast diversity of CAZymes present in the gut microbiota, highlighting the essential role of these microorganisms in digesting complex carbohydrates that the human genome alone cannot break down. This co-dependence between humans and their microbiota underscores the evolutionary importance of symbiosis, as humans have effectively outsourced much of their carbohydrate digestion to gut microbes.

CAZyme activity in the gut microbiome has several implications for health. Firstly, these enzymes release nutrients otherwise inaccessible to the host, enhancing digestive efficiency. Beyond nutrient release, the activity of CAZymes influences metabolic health, immune function, and even the risk of developing diseases such as obesity and diabetes. Moreover, the breakdown of complex carbohydrates by CAZymes is the initial step in their fermentation, where the chemical energy from carbon sources (complex carbohydrates) is converted into energy (ATP) that cells use in the anaerobic environment of the gut (Bernalier-Donadille, 2010; Grabitske & Slavin, 2008). CAZymes are classified into four families: Glycoside Hydrolases (GH) that cleave glycosidic bonds via hydrolysis, Polysaccharide Lyases (PL) that cleave glycosidic bonds through an elimination mechanism, Carbohydrate Esterases (CEs) that remove ester substituents from glycan chains and Glycosyltransferases (GTs) which are responsible for the assembly of complex carbohydrates from activated sugar donors. In addition, there are two other categories: Auxiliary Activities (AAs) that assist in the breakdown of polysaccharides, particularly through oxidative processes and, Carbohydrate-Binding Modules (CBMs) which facilitate the binding of CAZymes to specific carbohydrates, thereby enhancing their catalytic efficiency. Our study focuses on PLs and GHs, as they are responsible for the digestion of complex carbohydrates, a crucial process for human nutrition and health (Pan et al., 2021).

The primary end products of indigestible carbohydrate fermentation are short-chain fatty acids (SCFAs), including butyrate, acetate, and propionate, which serve as key energy substrates for the host, particularly for the intestinal epithelium (Simpson & Campbell, 2015; van der Hee & Wells, 2021). Bacterial fermentation substrates include indigestible carbohydrates (complex carbohydrates) derived from dietary fibers such as plant cell wall polysaccharides, resistant starch, soluble oligosaccharides, and endogenous products like mucin (Dalile et al., 2019). In the proximal colon, indigestible carbohydrates are fermented by saccharolytic bacteria producing SCFAs along with gases like CO2 and H2 (Topping & Clifton, 2001).

Following their production in the colon, SCFAs are rapidly absorbed by colonocytes, mainly via monocarboxylate transporter (MCT)-mediated active transport. Once absorbed, SCFAs enter the citric acid cycle in the mitochondria to generate ATP and provide energy to the cells (Schönfeld & Wojtczak, 2016). SCFAs that are not metabolized in the colonocytes are transported to the portal circulation. In humans, portal concentrations of SCFAs average 260 μM for acetate, 30 μM for propionate, and 30 μM for butyrate (Bloemen et al., 2009). In the liver, these SCFAs are utilized as energy substrates for hepatocytes, with acetate also serving as a precursor for cholesterol and fatty acid synthesis (Boets et al., 2017).

Beyond their role as an energy source for intestinal epithelial cells and in maintaining intestinal barrier function, SCFAs have emerged as critical regulators of immune function. Through mechanisms such as histone deacetylase (HDAC) inhibition and activation of G-protein-coupled receptors (GPCRs) like GPR41, GPR43, and GPR109A, SCFAs modulate both innate and adaptive immune responses (Liu et al., 2023). These mechanisms enable SCFAs to exert both pro-inflammatory and anti-inflammatory effects, depending on the cellular context and immune environment, highlighting their dual role in immune homeostasis (Kim, 2023).

In inflammatory bowel disease (IBD), encompassing ulcerative colitis (UC) and Crohn’s disease (CD), alterations in the gut microbiota are frequently accompanied by reduced SCFA production. This is attributed to a depletion of fiber-fermenting bacteria such as *Faecalibacterium prausnitzii* and *Roseburia intestinalis*. The resulting lower SCFA levels contribute to impaired energy metabolism in colonocytes, compromised barrier integrity, and reduced anti-inflammatory signaling, which collectively exacerbate chronic intestinal inflammation. Understanding the immunomodulatory functions of SCFAs and their disruption in IBD underscores their potential as therapeutic targets for restoring intestinal homeostasis and mitigating disease progression (Duncan et al., 2004; Sokol et al., 2008; Venegas et al., 2019).

Unique CAZyme profiles have been identified through comparisons between healthy controls and disease conditions, where these enzymes are underrepresented in cases of type 1 diabetes, colorectal cancer, and rheumatoid arthritis. Although certain carbohydrate-related functions are reduced, the overall CAZyme profile is largely preserved, maintaining significant metabolic functionality of carbohydrates even in disease states (Onyango et al., 2021). The development of techniques to characterize the CAZyme repertoire in individual microbiomes could open new avenues for treating disorders related to nutrition, metabolic dysfunction, intestinal mucosal barrier function, and other issues related to intestinal motility (Kaoutari et al., 2013).

The aim of this study is to characterize the gut CAZyme profile in different non-disease and disease cohorts from Argentina: A) a reference database controlled for healthy habits, B) a cohort of non-controlled participants from a wellness and prevention program (representing an average population living in industrialized cities), C) a cohort of patients with Inflammatory Bowel Disease (IBD) and D) a cohort of patients without Inflamatory Bowel Disease (control). By comparing these profiles, we seek to identify specific alterations in CAZyme composition associated with IBD, as well as the relationship between the CAZyme profile and SCFA production across the four groups. This research applies metagenomic sequencing and bioinformatics analysis to uncover functional differences in carbohydrate metabolism between healthy individuals and those with IBD. Our findings underscore the complexity of the non-linear relationship between complex carbohydrate metabolism and SCFA production. Additionally, the findings provide new insights into the role of gut microbiota in the pathogenesis of IBD.

## Materials and Methods

### Study cohorts

The Argentinian reference cohort was previously described (Rohr et al., 2023). Briefly, 200 volunteers were screened in four different cities of Argentina, after an extensive health and habits questionnaire and blood, urine and metabolome tests, 94 volunteers were selected for the reference-controlled program and stool samples were taken to analyze gut microbiome. This cohort is considered strictly adjusted for healthy habits (reference cohort). Another cohort of Fifty-four participants were randomly and anonymously selected from our wellness and prevention program. Participants agreed to participate anonymously in research studies upon accepted the terms and conditions of the program. This cohort was not strictly controlled for healthy habits and is considered a standard average population living in industrialized cities (average cohort). A third cohort is composed of thirty IBD patients that were recruited in the Cordoba Private Hospital in Argentina and donated stool samples to be analyzed in this project (Ethics protocol HP 4-285). Alongside this group, another **10** participants without IBD were selected from the same region and considered as a control group. This IBD and control cohort was further divided into IgG+ and IgG-subgroups. Separation of IgG+ or IgG-aliquots was performed as (Palm et al., 2014). 100 mg of frozen human fecal material was incubated in 1 ml Phosphate Buffered Saline (PBS) per 100 mg fecal material on ice for 1 hr. Fecal pellets were homogenized by manual pipetting and then centrifuged (50 x g, 15 min, 4C) to remove large particles. Fecal bacteria in the supernatants were removed (100 ul/sample), washed with 1 ml PBS containing 1% (w/v) Bovine Serum Albumin (BSA, Bovina Serum Albumin; staining buffer) and centrifuged for 5 min (8,000 x g, 4C) before resuspension in 1 ml staining buffer. A sample of this bacterial suspension (50 ul) was saved as the Pre-sort sample for sequencing analysis. After an additional wash, bacterial pellets were resuspended in 100 ml blocking buffer (staining buffer containing 20% Fetal Bovine Serum for human samples), incubated for 20 min on ice, and then stained with 100 ul staining buffer containing PE-conjugated Anti-Human IgG (1:100) for 30 min on ice. Samples were then washed 3 times with 1 ml staining buffer before cell separation.

Anti-IgG stained fecal bacteria were incubated in 1 ml staining buffer containing 50 ul Ultrapure Anti-PE Magnetic Activated Cell Sorting (MACS) beads (Miltenyi Biotec) (15 min at 4C), washed twice with 1 ml Staining Buffer (10,000 x g, 5 min, 4C), and then sorted by LS MACS separation columns (Miltenyi Biotech). After MACS separation, 500 ul of the negative fraction was collected for sequencing analysis (IgA negative fraction). For each sample, 500 ul of IgG-positive bacteria were collected, pelleted (10,000 x g, 5 min, 4C), and frozen along with the Pre-sort and IgG-negative samples at 80C for future use.

### Gut Microbiome Analysis

Stool samples from both cohorts were analyzed by low pass metagenomic sequencing of 1.3 million reads per sample following Illumina protocols on a NextSeq 550 sequencer. Bioinformatic analyses were performed using SqueezeMeta (Puente-Sánchez et al., 2020) and dbCAN3 software (Zheng et al., 2023) to identify CAZymes and microbiome functionality. To assign SCFAs (Short-Chain Fatty Acids) to the metagenomic samples, KEGG (Kyoto Encyclopedia of Genes and Genomes) identifiers were used. A total of 42 KEGG identifiers were employed to detect the enzymes involved in the metabolic pathway for SCFA production. SCFA enzyme abundance was calculated as FPM (Fragments per Million reads) to normalize the samples.

### Statistics Analysis

Data obtained from CAZymes and SCFA production was analyzed using *matplotlib* and *scipy* phyton libraries. Statistically significant was considered when p-values were > 0.05. Homoscedasticity of the samples was considered to decide whether to perform parametric or non-parametric tests.

## Results

### CAZymes Diversity in Gut Microbiomes: Reference-controlled vs. Average Cohorts

To characterize the microbiome composition in the Reference and Average cohorts, we analyzed genus-level alpha diversity (Fig 1A). The Reference cohort showed significantly higher genus-level alpha diversity compared to the Average cohort, indicating a more diverse microbial community at this taxonomic level.

**Figure 1.**
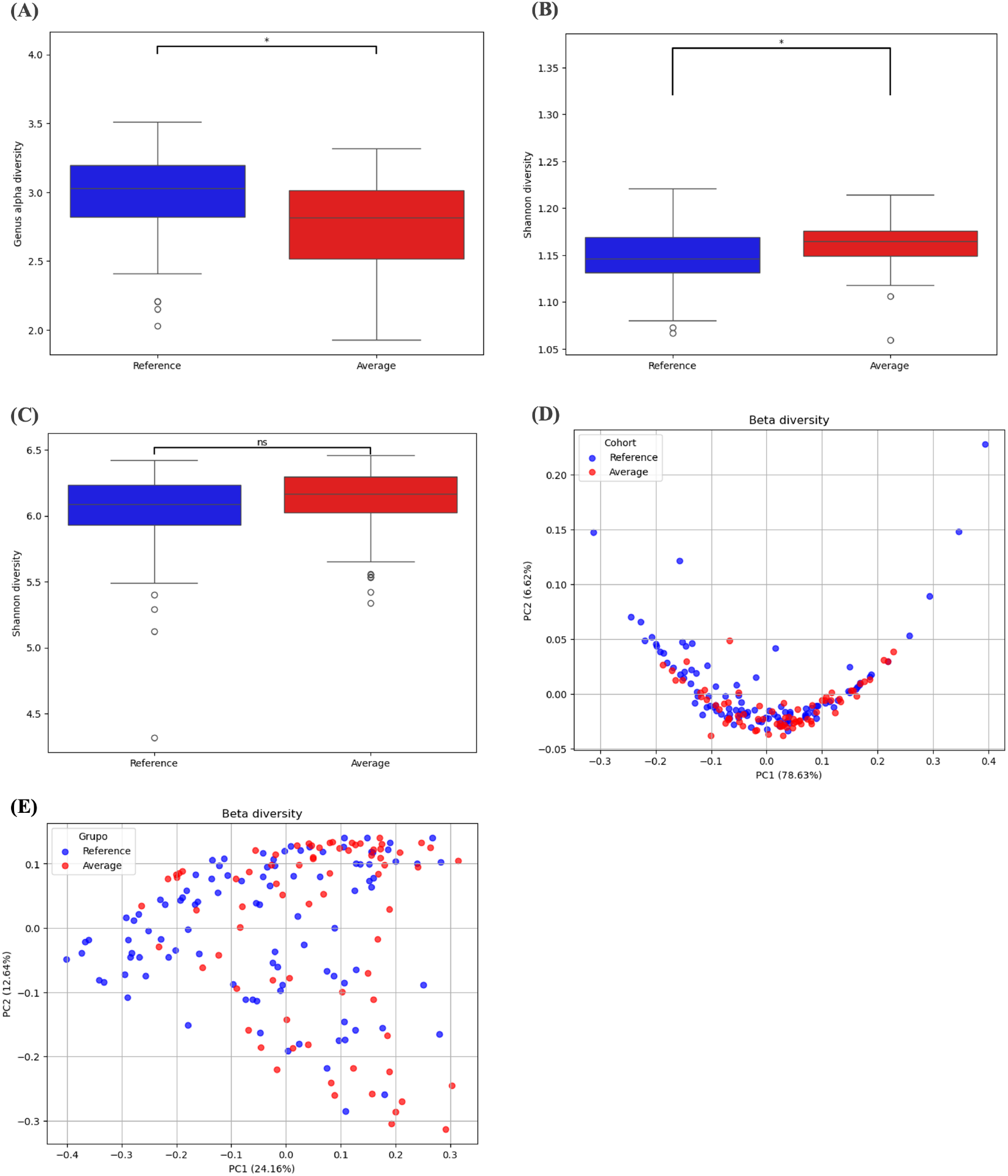
Diversity of bacteria and CAZymes families and subfamilies in gut microbiome of reference and Average cohort. **(A)** The boxplot illustrates genus-level alpha diversity, measured using Shannon diversity index, for individuals of the reference and Average cohort. A statistically significant difference between the groups is indicated (*p < 0.05). The median, interquartile range (IQR), and overall range of diversity values are shown for each cohort. **(B)** The boxplot displays Shannon diversity indices for alpha diversity of CAZyme families in average and reference cohorts (*p < 0.05, t-test). The median, interquartile range (IQR), and overall range of diversity values are shown for each cohort. **(C)** The boxplot displays Shannon diversity indices for alpha diversity of CAZyme subfamilies in average and reference groups (ns: non-significant, t-test). The median, interquartile range (IQR), and overall range of diversity values are shown for each cohort. **(D)** The PCoA plot shows beta diversity of CAZyme families with PC1 explaining 78.63% of the variance and PC2 explaining 6.62%. **(E)** The PCoA plot shows beta diversity of CAZyme subfamilies with PC1 explaining 24.16% of the variance and PC2 explaining 12.64% in CAZymes’subfamilies of both groups.

We then examined the distribution of CAZymes by calculating the abundance of CAZyme families across all samples. Interestingly, although genus-level alpha diversity was higher in the reference cohort, CAZyme alpha diversity at the family level was significantly greater in the Average cohort (Fig. 1B). However, no differences were observed in subfamily composition (Fig. 1C). Additionally, beta diversity analysis revealed comparable profiles for both families and subfamilies between the reference and Average cohorts, suggesting similar structural organization of CAZyme communities despite the observed differences in alpha diversity (Fig 1D and 1E).

Building on the observation that genus-level alpha diversity is significantly higher in the reference cohort while CAZyme alpha diversity at the family level is greater in the Average cohort, we aim to analyze in greater depth the potential differences in the functional composition of CAZyme families between the reference and Average cohorts. We analyzed the average abundance (FPM) of six major families: AA, CBM, CE, GH, GT, and PL (Supplementary figure 1). Significant differences were observed for two families: AA and PL. The AA family showed a significantly higher abundance in the Reference cohort compared to the Average cohort (p < 0.05; Supp. Fig. 1A). Conversely, the PL family exhibited a significantly higher abundance in the Average cohort (p < 0.05; Supp. Fig. 1F). For the remaining families (CBM, CE, GH, and GT), no significant differences were detected between the two cohorts, indicating comparable levels of abundance (Supp. Fig. 1B-E, respectively). These results indicate that while the AA and PL families exhibit significant variations in abundance between cohorts, the CBM, CE, GH, and GT families remain largely conserved. This suggests a stable functional profile for most CAZyme families, with only specific enzymatic groups undergoing notable shifts.

### SCFA production capacity and CAZymes abundance

To investigate the relationship between the metabolic capacity for short-chain fatty acid (SCFA) production and the abundance of carbohydrate-active enzymes (CAZymes) genes in the gut microbiome, we conducted a correlation analysis across both cohorts. In the Reference cohort and the Average cohort, we observed a statistically significant but weak negative correlation between these parameters (Fig. 2). Specifically, in the Reference cohort, the Spearman correlation coefficient was -0.17 (Fig. 2), indicating a slight decrease in SCFA production capacity with an increasing CAZyme abundance. Similarly, in the Average cohort, a Pearson correlation coefficient of -0.38 was observed (Fig. 2), showing a moderate negative relationship between these variables. These findings suggest an inverse relationship between CAZyme abundance and SCFA production capacity in the gut microbiome, albeit weak to moderate in strength. This negative correlation may indicate a trade-off between polysaccharide degradation and SCFA synthesis, potentially reflecting differences in microbial functional strategies across cohorts.

**Figure 2.**
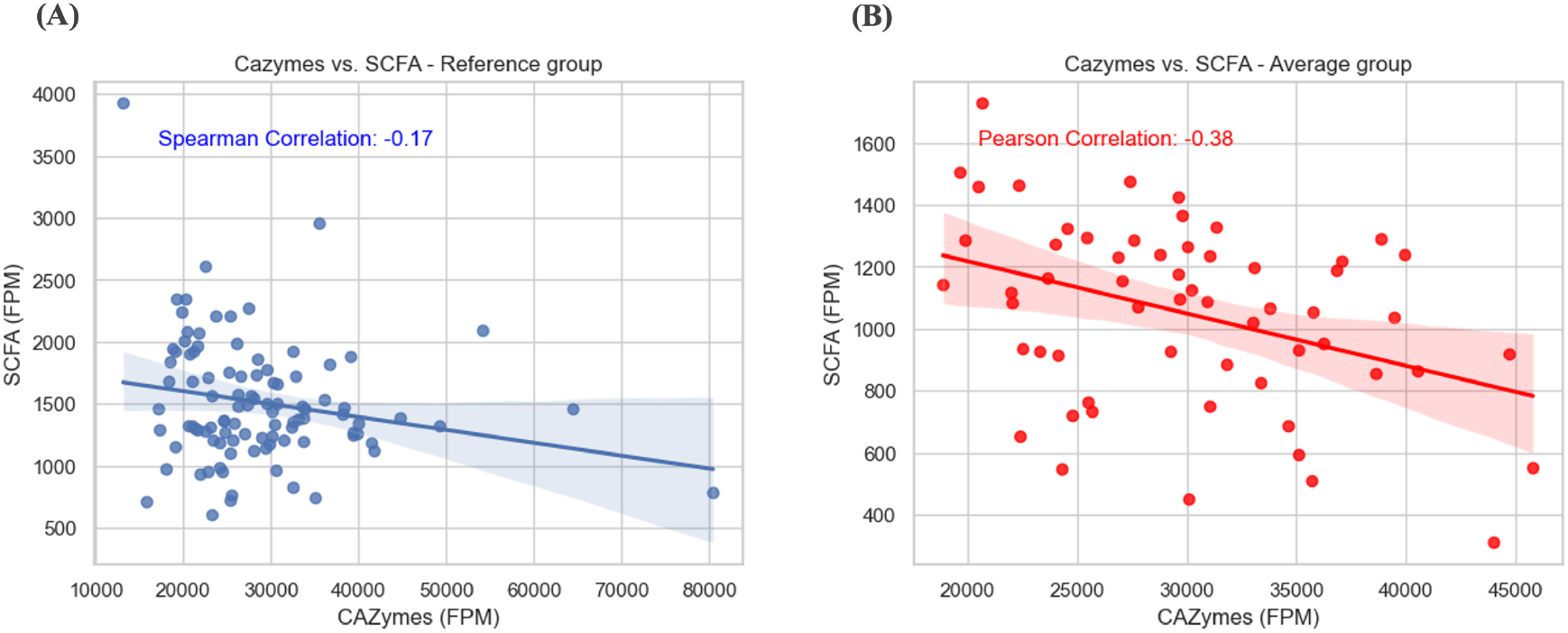
Relationship between CAZymes abundance and metabolic capacity of SCFA production. Scatter plots illustrate the relationship between CAZyme abundance (FPM) and SCFA production capacity (FPM) in the Reference **(A)** and Average **(B)** cohorts. Each point represents an individual sample. The blue regression lines indicate the linear trends, with shaded areas representing confidence intervals.

To further investigate the relationship between CAZyme abundance and SCFA metabolic capacity, we conducted an additional analysis focusing on the glycoside hydrolase (GH) and polysaccharide lyase (PL) families of CAZymes. These two families are well-known for their roles in breaking down complex carbohydrates and polysaccharides into simpler molecules, which serve as precursors for SCFA production in the gut microbiome. By examining these specific families, we aimed to gain deeper insights into how the enzymatic potential of the microbiome influences the metabolic pathways leading to SCFA synthesis.

The analysis of the correlation between SCFA production capacity and the abundance of the PL and GH families in both the Reference and Average cohorts reveals distinct patterns (Fig. 3). In the Reference group, a weak negative correlation was observed between PL abundance and SCFA metabolic capacity (Spearman correlation coefficient = -0.17, Fig. 3A), while GH abundance exhibited a slightly stronger negative correlation (Spearman correlation coefficient = -0.25, Fig. 3C). These results suggest that, in this cohort, higher abundance of these enzyme families is modestly associated with reduced SCFA production capacity.

**Figure 3.**
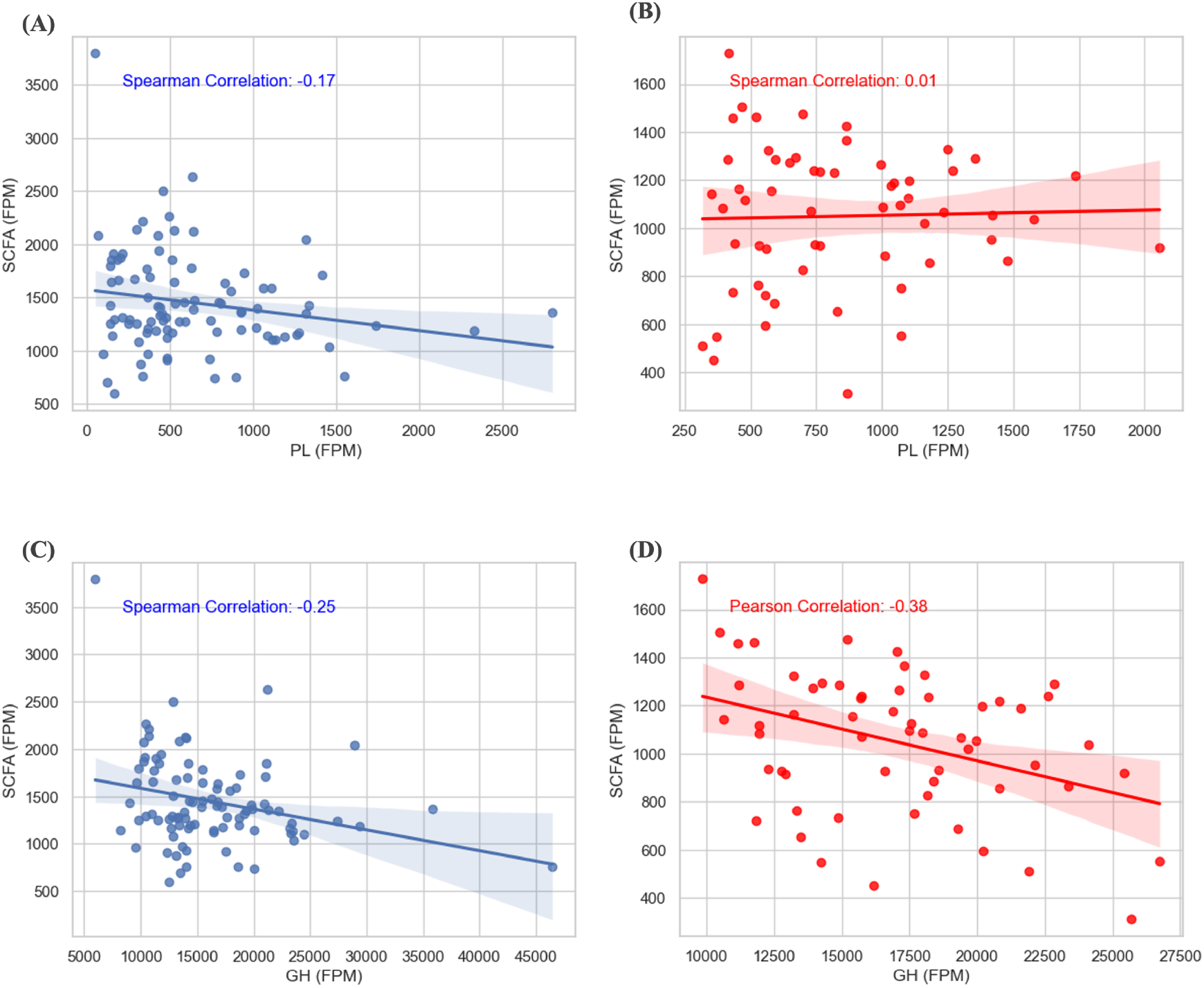
Relationship between GH and PL CAZymes families and metabolic capacity of SCFA. Scatter plots illustrate the relationship between PL and GH family abundance (FPM) and SCFA production capacity (FPM) in the Reference (**A and C**) and Average (**B and D**) cohorts. Each point represents an individual sample. The lines indicate the regression linear trends, with shaded areas representing confidence intervals.

In contrast, the Average cohort showed minimal correlation between PL abundance and SCFA production (Spearman correlation coefficient = 0.01, Fig. 3B), indicating little to no relationship. However, for GH abundance, a moderate negative correlation was observed (Pearson correlation coefficient = -0.38, Fig. 3D), suggesting a stronger association in this group compared to the reference cohort.

These findings highlight cohort-specific differences in the relationship between enzyme family abundance and SCFA metabolic capacity, with the GH family displaying a more consistent influence across both groups, while the role of the PL family appears to be less pronounced.

### Capabilities of the gut microbiome substrate degradation in the studied cohorts

Based on the available information about the CAZymes present in the samples and their corresponding substrate specificities, a targeted analysis was conducted focusing on two key enzyme families: PL and GH. These families were chosen because their substrate profiles include compounds that, during fermentation, are metabolized into short-chain fatty acids (SCFAs) as end products. Notably, some known SCFA precursors were not detected in the dbCAN3 analysis. This absence could be attributed either to (1) the lack of corresponding enzyme annotations in the dbCAN3 database or (2) the absence of these enzymes in the analyzed samples. The undetected substrates included arabinoxylan, lactose, inulin, milk oligosaccharides, gum arabic, guar gum, laminaria, and stachyose. Conversely, substrates identified as being degraded by the CAZymes present in the samples’ microbiota included resistant starch, cellulose, xylan, pectin, fructo-oligosaccharides, beta-glucans, and raffinose. These substrates formed the basis for a detailed analysis of SCFA production potential.

In Supplementary Figure 2, the detected substrate degradation capabilities of PL and GH families are represented. The abundance is quantified as the average fragments per million reads (FPM) of specific enzymes for each substrate. As observed, the average cohort exhibits a significantly greater degradative capability for cellulose (Suppl. Fig. 2C) and pectin (Suppl. Fig. 2F), while the reference cohort shows higher degradative capability for raffinose (Suppl. Fig. 2G). Both the reference and average cohorts display a high abundance of enzymes responsible for xylan degradation (Suppl. Fig. 2H), followed by those for beta-glucans (Suppl. Fig. 2B). In both groups, the least abundant enzymes are those responsible for raffinose degradation (Suppl. Fig. 2G).

The substrates analyzed are known precursors of SCFAs, making their degradation capabilities essential for understanding SCFA synthesis in the gut microbiome. To explore this relationship, we examined the correlations between the degradative capacities of these substrates and the SCFA production potential across individual samples within each cohort.

Our findings revealed significant negative correlations for all substrates except raffinose, suggesting that higher degradation capabilities for most substrates are associated with lower SCFA production potential (Fig. 4). This trend was evident across both cohorts, with stronger negative correlations consistently observed in the Average cohort compared to the Reference cohort.

**Figure 4.**
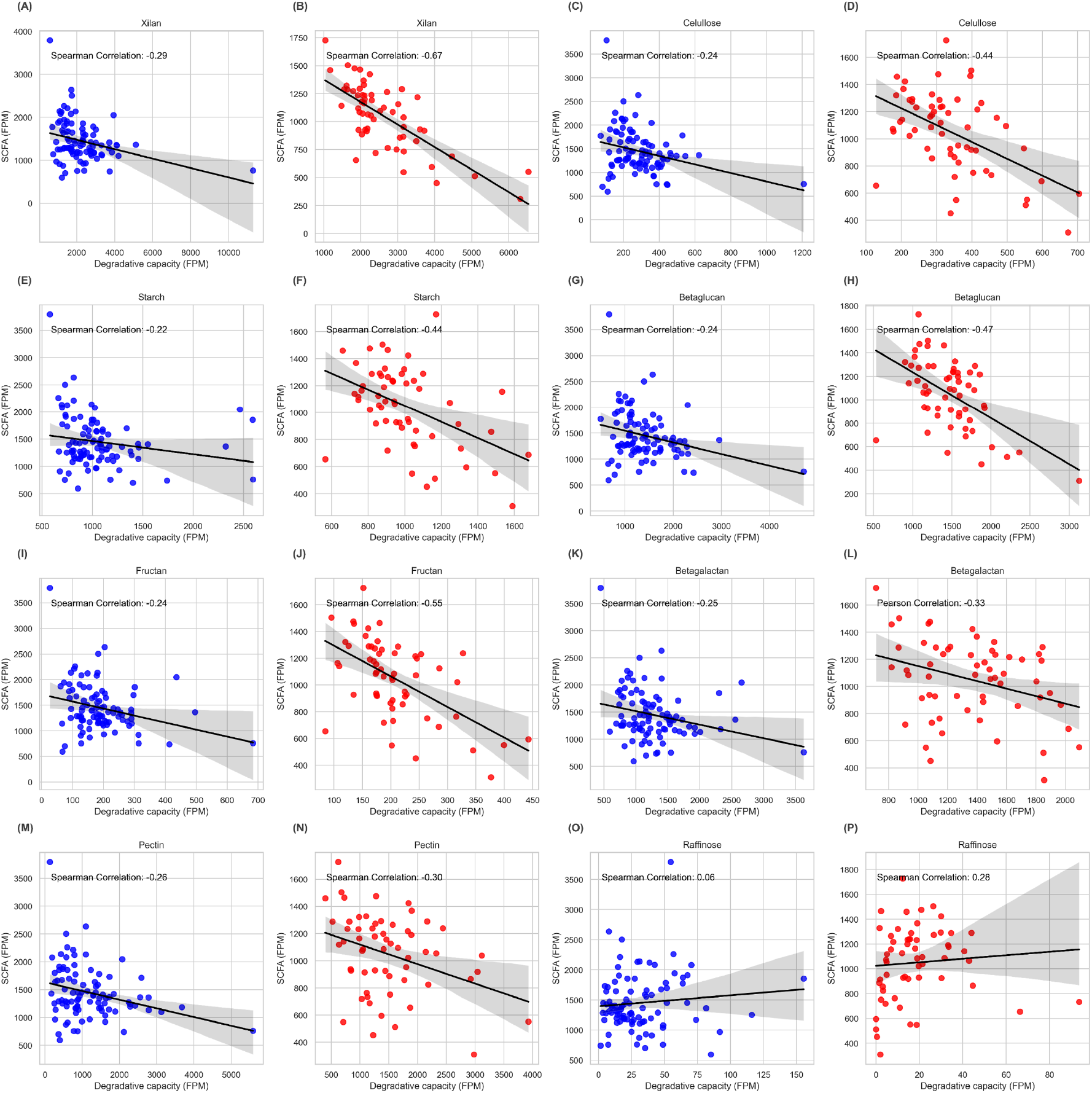
Relationship between SCFA-related substrate degradation capacity by CAZymes and metabolic capacity of SCFA production. Scatter plots illustrate the relationship between substrate degradation capacity (FPM) and SCFA production capacity (FPM) including Xylan (A-B), Cellulose (C-D), Starch (E-F), Betaglucan (G-H), FOS (I-J), Galactan (K-L), Pectin (M-N), Raffinose (O-P), in the Reference (blue) and Average (red) cohorts. Each point represents an individual sample. The black regression lines indicate the linear trends, with shaded areas representing confidence intervals.

For instance, Xylan showed a correlation of -0.29 in the reference cohort (Fig. 4A) but a stronger negative correlation of -0.57 in the Average cohort (Fig. 4B). Similarly, cellulose displayed a shift from -0.24 in the reference cohort (Fig. 4C) to -0.44 in the Average cohort (Fig. 4D). Beta-glucans followed the same pattern, with correlations of -0.25 and -0.47 for the reference (Fig. 4G) and Average cohorts (Fig. 4H), respectively. This trend was also observed for starch (-0.24 in the reference group vs. -0.44 in the Average group, Fig 4E-F), fructans (-0.24 in the reference vs. - 0.55 in the Average cohort, Fig. 4I-J) and pectin (-0.26 in the reference vs. -0.30 in the Average cohort, Fig. 4M-N).

Interestingly, raffinose was an outlier, showing no significant negative correlation in either cohort (Fig. 4O-P). In fact, in the Average cohort, it presented a weak positive correlation (0.28, Fig P), potentially reflecting different microbial or enzymatic dynamics in this cohort.

### Cazymes in inflammatory bowel disease

With the aim of evaluating the status of CAZymes in two different conditions of Inflammatory Bowel Disease (IBD cohort), patients were recruited with Ulcerative Colitis (UC) and Crohn disease (CD), as well as people without IBD (control cohort) at Hospital Privado Universitario de Cordoba. Since we did not observe any differences between UC and CD group in either genus or Cazymes composition, we show here both cohorts in a unique one named as IBD cohort. To begin with, we analyzed genus diversity in IBD cohort compared to control cohort (Fig. 5A). IBD samples had a significantly lower genus alpha diversity than control samples (p= 0.03).

**Figure 5.**
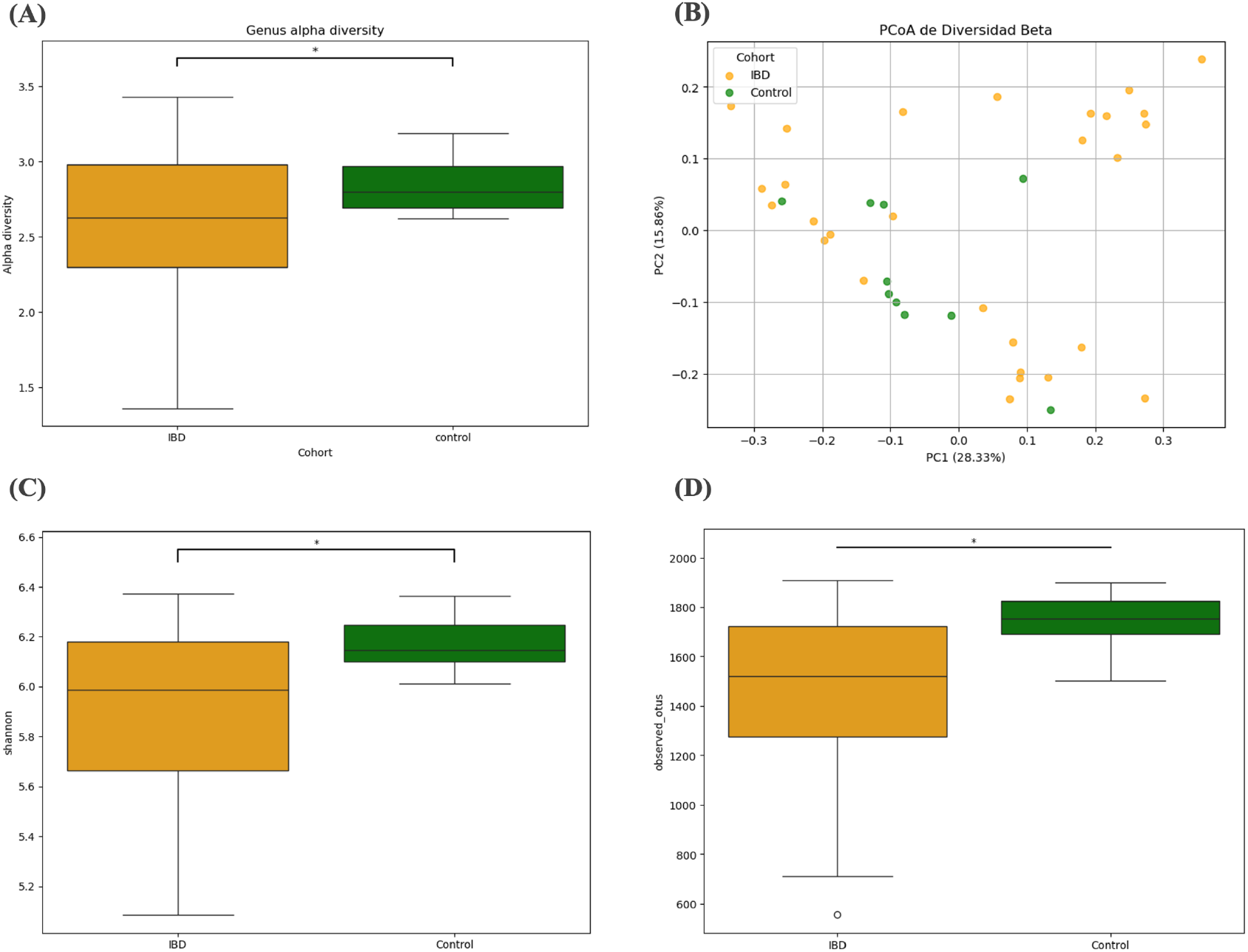
Comparison of genus-level alpha diversity and CAZymes subfamilies alpha and beta diversity between IBD and control cohorts. **(A)** The boxplot illustrates genus-level alpha diversity, measured using Shannon diversity index, for individuals with inflammatory bowel disease (IBD, orange) and control (green) subjects. A statistically significant difference between the groups is indicated (*p < 0.05). The median, interquartile range (IQR), and overall range of diversity values are shown for each cohort. **(B)** PCoA plot shows beta diversity of CAZyme subfamilies in IBD (orange) and control (green) groups with PC1 explaining 28.33% of the variance and PC2 explaining 15.59%. **(C)** The boxplot displays Shannon diversity indices for alpha diversity of CAZyme subfamilies in IBD (orange) and control (green) groups (*p < 0.05, t-test). **(D)** The boxplot displays observed OTUs of CAZyme subfamilies in IBD (orange) and control (green) groups (*p < 0.05, t-test).

Then, we explore CAZymes subfamily’s diversity in both cohorts. The beta and alpha diversity of CAZyme subfamilies were analyzed in individuals with IBD (IBD group) and without IBD (control group). The PCoA plot (Fig. 5B) illustrates the beta diversity, showing a partial separation between the IBD (orange) and control (green) groups along the first principal coordinate (PC1), which explains 28.33% of the variance, and the second principal coordinate (PC2), which explains 15.59%. This indicates differences in the CAZyme subfamily composition of the gut microbiome between the groups. The boxplot (Fig. 5C) depicts the Shannon diversity index for alpha diversity. A t-test revealed a statistically significant difference (*p < 0.05) in Shannon diversity, with the control group exhibiting higher diversity compared to the IBD group. Additionally, the observed OTUs boxplot (Fig. 5D) demonstrates a significant reduction in the richness of CAZyme subfamilies in the IBD group compared to the control group (*p < 0.05, t-test). These results suggest that inflammatory bowel disease is associated with both reduced functional diversity and distinct compositional changes in the gut microbiome’s CAZyme subfamilies, highlighting a potential disruption in enzymatic functions linked to the disease.

To further investigate the functional composition of the gut microbiome in IBD, the samples from the IBD group were separated into two subsets: IgG-positive (pos) and IgG-negative (neg) fractions. The IgG-positive samples represent bacteria that were recognized and bound by immunoglobulin G (IgG), indicating a potential immune response targeting these taxa. In contrast, the IgG-negative samples include bacteria that were not recognized by IgG, suggesting they are less likely to be involved in direct host immune interactions. This separation allows for a deeper exploration of the functional and compositional differences between immune-targeted and non-targeted microbial populations within the gut microbiome of IBD patients, shedding light on the interplay between microbial functionality and host immune responses.

The PCoA plot of beta diversity (Fig. 6A) shows a distinct clustering of the control and group IBD subgroups, with clear separation between IgG-positive and IgG-negative samples along the first principal coordinate (PC1, 35.18%of variance). This indicates that the microbial composition differs significantly across these groups. The alpha diversity analyses further support these findings. The Shannon index (Fig 6B) reveals a significant decrease in diversity for the IgG-positive samples compared to both IgG-negative and reference samples (p < 0.0001). Similarly, the observed OTUs follow the same trend, with IgG-positive samples showing the lowest richness (Fig 6C). These results suggest that bacteria recognized by IgG exhibit reduced diversity and richness in CAZymes subfamilies, potentially reflecting an immune-driven selection process in IBD. Conversely, the IgG-negative fraction retains greater diversity, closer to that of the reference group, highlighting the differential impact of immune recognition on the microbiome composition in IBD.

**Figure 6.**
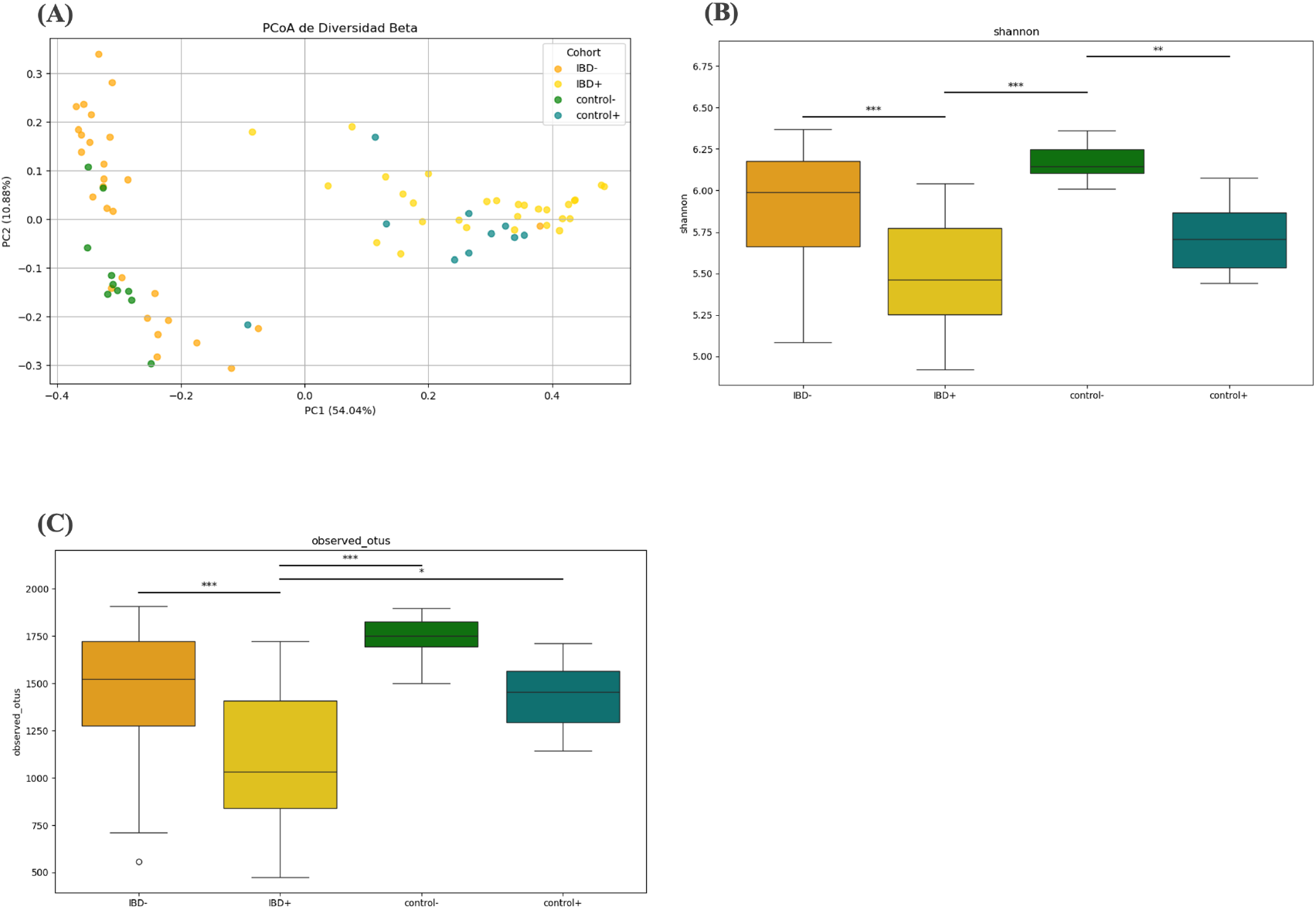
Beta and Alpha Diversity of CAZyme Subfamilies in IgG-Positive and IgG-Negative Samples from Both Control and IBD Patients. Beta and alpha diversity analyses of CAZyme subfamilies in IgG-negative control (control-), IgG-positive control (control+), IgG-negative IBD (IBD-) and IgG-positive IBD (IBD+) samples. **(A)** The PCoA plot illustrates beta diversity, with the first principal coordinate (PC1) explaining 54.04% of the variance. Boxplots compare Shannon diversity **(B)** and observed OTUs **(C)** across groups. ANOVA p<0.05. Red numbers indicate Post Hoc analysis results.

Finally, with the aim of thoroughly evaluating the role of CAZymes in IBD, we analyzed the abundance of subfamilies previously reported to be associated with IBD (Labourel et al., 2023). We then analyzed GH2 (Fig. 7A), GH16 (Fig. 7B), GH29 (Fig. 7C), GH31 (Fig. 7D), GH33 (Fig. 7E), GH35 (Fig. 7F), GH95 (Fig. 7G), GH98 (Fig. 7H), GH109 (Fig. 7I), GH112 (Fig. 7J) and GH136 (Fig. 7K). The analysis revealed no significant differences between the control and IBD groups (Fig 7). However, significant differences were observed between IgG-positive (IBD+ and control +) and IgG-negative (IBD- and control-) subgroups. Almost all studied enzymes were more prevalent in IgG-negative samples except for GH31 (Fig. 7D), GH35 (Fig. 7F) y GH109 (Fig. 7I) which were more prevalent in IgG-positive samples.

**Figure 7.**
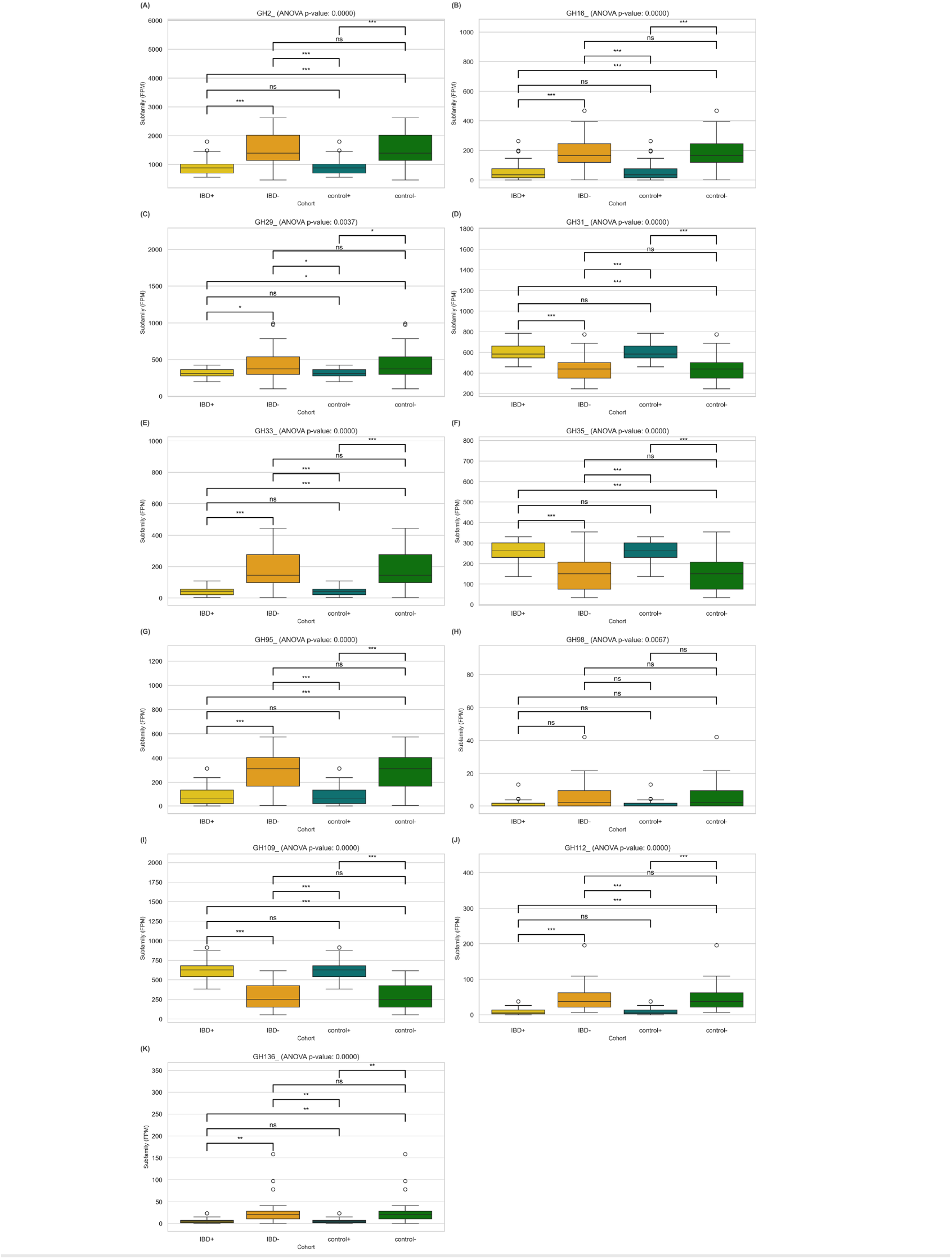
Subfamily abundance levels of glycoside hydrolases across different cohorts. Boxplots show the abundance levels of fragments per million (FPM) of glycoside hydrolase subfamilies GH2 **(A)**, GH16 **(B)**, GH29 **(C)**, GH31 **(D)**, GH33 **(E)**, GH35 **(F)**, GH95 **(G)**, GH98 **(H)**, GH109 **(I)**, GH112 **(J)** and GH136 **(K)** in IBD-positive (IBD+), IBD-negative (IBD−), control-positive (control+), and control-negative (control−) groups. Statistical significance was determined using ANOVA with post-hoc comparisons: *p < 0.05, **p < 0.01, ***p < 0.001, ns = not significant.

## Discussion

This study aimed to characterize the profile and functionality of carbohydrate-active enzymes (CAZymes) in the gut microbiome through metagenomic analysis of gut microbiota from two different cohorts: a reference-controlled population of healthy habits and individuals participating in a chronic disease prevention program that were not controlled for healthy habits (Average cohort). Additionally, the composition and diversity of CAZymes were examined in the context of Inflammatory Bowel Disease (IBD) to evaluate the profiles of CAzymes under disease conditions compared to healthy controls. Interestingly, the findings revealed that the CAZyme profile remains largely stable across different health conditions, including IBD. This suggests a remarkable functional resilience of the gut microbiome in maintaining carbohydrate metabolism, even in the presence of chronic diseases, highlighting the conserved nature of these enzymatic pathways. These results contribute to a deeper understanding of the gut microbiome’s role in health and disease and provide a foundation for future studies exploring its therapeutic potential. In our analysis, the Average cohort showed greater CAZyme alpha diversity at the family level, despite no significant differences in subfamily composition. Beta diversity analysis revealed comparable structural organization of CAZyme communities between the two cohorts, suggesting that variations in microbial diversity do not necessarily translate into major differences in the functional organization of CAZymes. These results confirm the findings of Onyango SO and his colleagues, who observed that the functionality of CAZymes is conserved across different populations (Onyango et al., 2021).

To investigate the relationship between the abundance of CAZymes and SCFA production capacity, we analyzed the correlations between these two indicators. We observed that the abundance of certain CAZymes is largely inversely proportional to SCFA production capacity. Specifically, in both datasets, the correlations were negative and below 0.5, indicating that there is no direct relationship between the abundance of CAZymes and the capacity to produce SCFAs. Given the diverse functions of CAZyme families, we proceeded to analyze the relationship between CAZyme families involved in the degradation of complex carbohydrates, specifically GHs and PLs, and SCFA production capacity. Again, both factors showed low Spearman correlations with SCFA levels. This suggests that the degradative potential of these enzymes does not directly translate into greater SCFA production. An explanation for the low correlation values observed between the abundance of CAZymes and SCFA production is that complex carbohydrates (CCs) can follow various metabolic pathways beyond fermentation into SCFAs. For instance, CCs like glycogen can be channeled into glycolysis or the hexosamine biosynthetic pathway, leading to the production of glycoproteins and glycolipids that play essential roles in cellular signaling and membrane structure (Ibrahim & Anishetty, 2012). Additionally, CAZymes are not solely involved in the breakdown of complex carbohydrates. They also play a crucial role in the degradation of N-glycans, which are common post-translational modifications of proteins on asparagine residues. These enzymes act on N-glycans produced endogenously or derived from dietary sources, providing essential nutrients for the intestinal microbiota. The primary families of enzymes involved in this function include GH18, GH85, and PNGases, which hydrolyze specific bonds in the core pentasaccharide structure of N-glycans (Crouch et al., 2022).

Based on these observations, it could be suggested that increasing complex carbohydrates in the diet does not necessarily result in an increase of SCFAs such as butyrate. This has important implications when planning dietary patterns in people who have low concentrations of SCFAs in their intestines or low metabolic capacity of SCFA pathways in their gut microbiomes. Therefore, the dietary pattern in these cases should be rethought including other variables.

A previous study compared the abundance and distribution of 220 recovered CAZyme families in saliva and stool samples from patients with colorectal cancer, rheumatoid arthritis, and type 1 diabetes with those of healthy subjects. The multivariate discriminant analysis suggested that the disease phenotype did not significantly alter the CAZyme profile, indicating a functional conservation of carbohydrate metabolism in these disease states (Onyango et al., 2021). In contrast, in IBD patients, we observed both a reduction in functional diversity and distinct compositional changes in CAZyme subfamilies of gut microbiome of this patients compared with healthy controls, suggesting a disruption in enzymatic functions associated with the disease. Notably, genes encoding mucin-degrading CAZymes, previously reported to be more prevalent in the microbiome of IBD patients (Labourel et al., 2023), were also analyzed in our IBD and control groups. However, we did not observe significant differences in the prevalence of these genes between the two groups. On the other hand, differences were identified between igG positive and igG negative subgroups in both control and IBD gut microbiomes.

The observed differences in CAZyme abundance between IgG-positive and IgG-negative bacterial populations suggest that the immune system selectively targets specific bacteria that may have inflammatory functions or influence the metabolic activity of the microbiome environment (Vujkovic-Cvijin et al., 2022). Notably, only three CAZyme families—**GH95, GH29, and GH109**—were found to be significantly more abundant in IgG-positive samples. These enzymes are known for their roles in mucin degradation and oligosaccharide processing, which could facilitate bacterial interactions with the host mucosal barrier and subsequent immune recognition. The higher abundance of **GH95** (α-fucosidase) in IgG-positive samples may indicate the presence of bacterial taxa that specialize in degrading fucosylated mucins, potentially exposing the underlying epithelial surface to immune detection. Similarly, **GH29** (another α-fucosidase) and **GH109** (α-N-acetylgalactosaminidase) could further contribute to the breakdown of specific mucin glycans, creating conditions that favor bacterial adherence and antigen exposure. These processes might trigger IgG-mediated immune responses, as the host recognizes these bacteria as potentially harmful due to their capacity to degrade protective mucosal barriers.

These findings suggest that the immune system does not uniformly target all mucin-degrading bacteria but rather focuses on specific taxa or activities, as reflected by the enrichment of only a subset of CAZyme families in IgG-positive samples. This selective recognition may have significant implications for the composition and function of the microbiome, particularly in IBD patients, where mucosal barrier integrity is compromised.

Further studies are needed to determine the specific bacterial taxa associated with **GH95, GH29, and GH109** enrichment in IgG-positive samples and their potential contributions to mucosal barrier disruption or immune activation. Insights into these interactions could provide a deeper understanding of how selective immune targeting shapes microbiome composition and function in both healthy and disease states.

These findings highlight the multifaceted roles of CAZymes in carbohydrate metabolism and suggest that their abundance alone may not directly predict SCFA production, as carbohydrates can be diverted to alternative pathways critical for host and microbial functions. Moreover, subfamilies diversity and composition were altered in gut microbiome of IBD patients. However, we did not find any specific subfamily altered in this group, such as the previously reported mucin degrading CAZyme.

The lack of a direct relationship between CAZyme abundance and SCFA production highlights the need for a more nuanced approach to dietary planning. Diets designed to enhance SCFA production should account for the metabolic versatility of carbohydrates and their alternative fates within the gut ecosystem. Integrating microbiome functional profiling with dietary interventions could offer a more personalized strategy for improving gut health. Further studies focusing on the interplay between these pathways and their metabolic outcomes could provide deeper insights into the complex relationship between dietary intake, microbial activity, and host health.

## Acknowledgements

The authors want to thank all members of the lab and the Heritas team for helpful discussions during the preparation of this manuscript.

## Data and Code availability

https://github.com/militrot/cazymes. Dataset availability upon request after publication.

## Supplemental Materials

**Supplementary figure 1.**
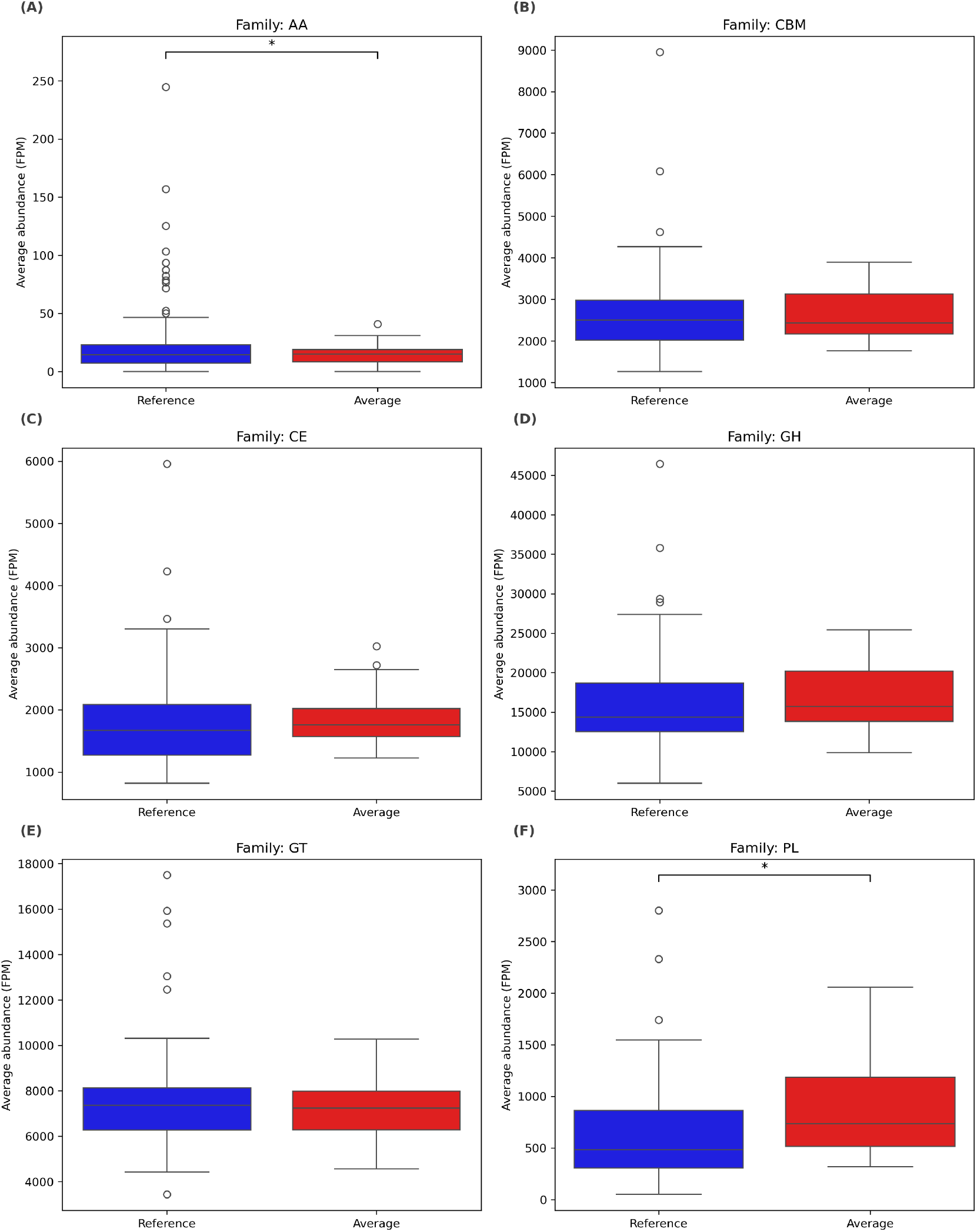
Comparison of CAZyme family abundance between Reference and Average cohorts. Boxplots show the average abundance (FPM) of six CAZyme families AA **(A)**, CBM **(B)**, CE **(C)**, GH **(D)**, GT **(E)**, and PL **(F)** in the Reference (blue) and Average (red) cohorts. A statistically significant difference between the groups is indicated (*p < 0.05). The median, interquartile range (IQR), and overall range of diversity values are shown for each cohort.

**Supplementary figure 2.**
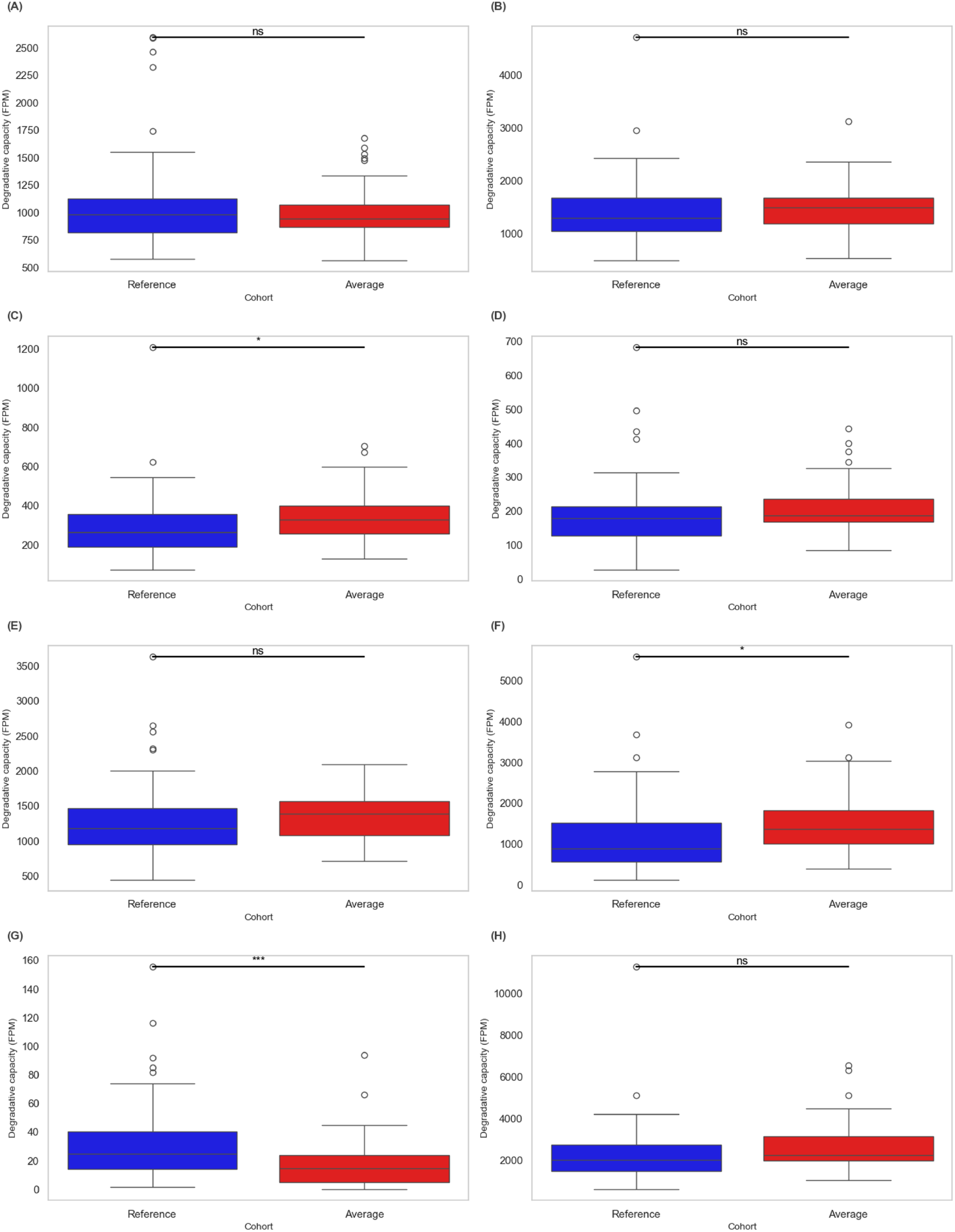
SCFA-substrate degradation capacity by CAZymes. Boxplots illustrate the degradation capacity of SCFA substrates, including Starch (A), Betaglucan (B), Cellulose (C), FOS (D), Galactan (E), Pectin (F), Raffinose (G), and Xylan (H), measured as fragments per million reads (FPM), for individuals in the Reference (blue) and Average (red) cohorts. Statistically significant differences between the groups are indicated (*p < 0.05, ***p < 0.001). The median, interquartile range (IQR), and overall range of degradation capacity values are shown for each cohort.

